# COPS5 maintains spermatogenesis gene expression and piRNA-mediated transposon silencing

**DOI:** 10.64898/2026.11.12.766086

**Authors:** Jiaolong Huang, Qian Huang, Cheng Zheng, Yunan Li, Dylan Millikin, Grace M. Swanson, Yi Tian Yap, Wei Li, Chen Chen, Stephen A. Krawetz, Yue Chen, Shuo Shi, Ling Zhang, Zhibing Zhang

## Abstract

COPS5 (CSN5/JAB1), a multifunctional subunit of the COP9 signalosome, is required for male germ cell survival, but how it acts is unclear. We combined a germ cell-specific *Cops5* conditional knockout (cKO) mouse model with transcriptomic, proteomic and small RNA analysis of P16 testes. *Cops5* loss caused progressive spermatogenic arrest after postnatal day 16. At both the mRNA and protein levels, 59 genes were downregulated, including piRNA biogenesis factors such as PIWIL1/MIWI, PIWIL2/MILI, Tudor-domain proteins, DDX4, MAEL and MOV10L1. In *Cops5* cKO testes, pachytene piRNAs decreased in abundance and diversity, falling from about 56% to 18–30% of total small RNAs, while the 1U bias was retained. Consistent with this, the transposable elements LINE1 and IAP were activated. In human testes, COPS5 was high in spermatogenic cells but reduced by 56.4% in the germ cells of patients with non-obstructive azoospermia (NOA). Thus, COPS5 maintains piRNA homeostasis and transposon silencing, and its loss may contribute to human spermatogenic failure.

## 1. Introduction

Spermatogenesis is the process by which diploid spermatogonia develop into spermatozoa, and it takes place in the seminiferous epithelium of the testis. It proceeds through three phases: the mitotic proliferation of spermatogonia, the meiotic division of spermatocytes, and spermiogenesis, in which haploid round spermatids undergo a marked morphological change into elongated sperm^1^. This process depends not only on germ cells but also on the somatic cells of the testis, including Sertoli cells, Leydig cells, and peritubular myoid cells^2^. Defects in this process reduce sperm production and, in severe cases, cause azoospermia, the absence of sperm in the ejaculate^3,4^.

Spermatogenesis is precisely regulated, and a number of key regulators, including COPS5 (COP9 signalosome subunit 5), have been identified. COPS5, also known as CSN5 or JAB1, is the catalytic subunit of the COP9 signalosome (CSN), a conserved complex consisting of eight subunits^5–8^. COPS5 regulates the stability of various proteins and is involved in cell cycle progression, the DNA damage response, development, and tumorigenesis ^9^. COPS5 was originally identified as JAB1 (Jun activation domain-binding protein 1), a coactivator that interacts with transcription factors such as c-Jun and can thus modulate gene transcription^10^. COPS5 is expressed in most tissues but is particularly abundant in the testis^11^. Our previous studies have shown that COPS5 is required in several cell types of the testis. Mice with germ cell-specific deletion of *Cops5* have markedly reduced sperm numbers, show germ cell apoptosis and defects in acrosome biogenesis, and are infertile^11^. On one hand, germ cell-specific deletion of *Cops5* markedly reduces sperm number, exhibiting defects in acrosome biogenesis, germ cell apoptosis and are infertile^11^. In other systems, like the deletion of *Cops5* in testicular smooth muscle cells causes growth retardation, abnormal development of the reproductive organs, and impaired endocrine function^12^. Sertoli cell-specific deletion of *Cops5* disrupts Sertoli cell polarity and the blood–testis barrier and leads to progressive loss of germ cells^2^. On the other hand, how COPS5 mediates the survival of male reproductive germ cells remains to be defined.

During germ cell development, the genome is also at risk from transposable elements (TEs) ^13^, and uses a PIWI-based defensive system to keep them at bay. Failure results in meiotic arrest and male infertility. In animals, this defense is mainly mediated by PIWI-interacting RNAs (piRNAs), a class of small non-coding RNAs of about 24 to 32 nucleotides that are expressed almost exclusively in the germline and that bind to PIWI proteins^14,15^. In mice, piRNAs are produced in two temporally distinct waves. In fetal prospermatogonia, piRNAs bound to MILI (PIWIL2) and MIWI2 (PIWIL4) guide de novo DNA methylation of TE regulatory regions through the DNMT3L–DNMT3A complex and thereby establish transcriptional silencing^16–19^. In postnatal germ cells, a second wave of piRNAs, known as pachytene piRNAs, is loaded mainly onto MILI and MIWI (PIWIL1) and acts largely at the post-transcriptional level^20^. Genetic studies in mice have shown that the piRNA pathway is essential for spermatogenesis: male mice lacking *Miwi*, *Mili*, *Miwi2*, *Mael*, *Tdrd1*, *Tdrd5*, *Tdrd9*, *Mov10l1*, *Tdrkh*, *Pnldc1*, *Henmt1*, or *Tex15* are sterile and show increased TE expression together with defects in meiotic progression^21–32^. Consistent with these findings, loss-of-function variants in human piRNA pathway genes such as PNLDC1 and TDRKH cause non-obstructive azoospermia (NOA) with meiotic or spermatogenic arrest^33,34^. These observations show that piRNA biogenesis is required for germ cell development and that its failure is a direct cause of human spermatogenic failure.

To define how COPS5 regulates germ cell development, germ-cell specific knockout mice were subjected to RNA sequencing, proteomics, and small RNA sequencing. These analyses allowed us to identify the genetic programs that depend on COPS5 in the developing testis. We show that the piRNA pathway is one of the major mechanisms by which COPS5 controls germ cell survival and spermatogenesis.

## 2. Materials and methods

### 2.1 Ethics statement

All animal experiments were conducted in accordance with institutional guidelines and approved by Wayne State University Institutional Animal Care with the Program Advisory Committee (Protocol number: 24-02-6561).

### 2.2 Generation of conditional *Cops5* knockout mice

Mice were maintained on a C57BL/6J background. Using our previously published method, we generated germ cell conditional knockout mice by crossing *Cops5 ^flox/flox^* females with *Stra8-iCre* males to obtain *Cops5* conditional knockout (*Cops5* cKO) mice^11^. Littermates lacking the Cre transgene were used as controls. Successful inactivation of *Cops5* in germ cells was verified by genotyping of the floxed allele and by Western blot analysis of testicular lysates.

### 2.3 Western blot analysis

Testes were homogenized in RIPA lysis buffer supplemented with protease and phosphatase inhibitors. Lysates were clarified by centrifugation, and protein concentrations were determined using a BCA assay. Equal amounts of protein were separated by SDS– PAGE and transferred onto PVDF membranes. Membranes were blocked with 5% non-fat milk in TBS–Tween 20 and incubated overnight at 4 °C with primary antibodies diluted. After three washes with TBS–Tween 20, membranes were incubated with HRP-conjugated secondary antibodies. Signals were detected using enhanced chemiluminescence on a ChemiDoc imaging system and quantified with ImageJ using densitometry values normalized to the corresponding loading control.

#### 2.3.1 Subcellular fractionation

Testicular lysates from adult mice were fractionated into cytoplasmic and nuclear components using a commercial fractionation kit according to the manufacturer’s protocol. Briefly, testes were homogenized in ice-cold lysis buffer, nuclei were pelleted by low-speed centrifugation, and the resulting nuclear pellet and cytoplasmic supernatant were separated further by high-speed centrifugation. The two fractions were mixed with SDS– PAGE sample buffer and analyzed by Western blot analysis as described above. α-Tubulin and lamin B were used as markers for the cytoplasmic and nuclear fractions, respectively.

### 2.4 Histology on tissue sections

Testes were fixed overnight in 4% paraformaldehyde, dehydrated through a graded ethanol series, embedded in paraffin, and sectioned at 5 μm thickness. Sections from at least five mice per group were stained with hematoxylin and eosin (H&E) according to standard protocols. Spermatogenic stages were assigned by examining at least 200 seminiferous tubules per mouse across five systematically sampled fields. Seminiferous tubule morphology, germ cell composition, and spermatogenic stages were examined under a light microscope (Olympus BX51, USA), and photographs were taken with a camera (Jenoptik Laser, Prog Res C14, Germany).

### 2.5 RNA-seq

Total RNA was extracted from 16-day-old testes using TRIzol reagent according to the manufacturer’s instructions. The quality of total RNA was assessed using an Agilent Fragment Analyzer (Agilent Technologies, Santa Clara, CA), and only RNA with RQN > 7 was used for subsequent mRNA-seq library preparation and sequencing. Approximately 500 ng of total RNA was used for stranded mRNA-seq library preparation by following the NEB Directional mRNA-seq sample preparation guide (New England Biolabs, Ipswich, MA). The first step in the workflow involved purifying the poly-A containing mRNA molecules using poly-T oligo-attached magnetic beads. Following purification, the mRNA was fragmented into small pieces using divalent cations under elevated temperature. The cleaved RNA fragments were copied into first strand cDNA using reverse transcriptase and random primers. These cDNA fragments then went through an end repair process, the addition of a single ‘A’ base, and then ligation of the adapters. The products were then purified and enriched with PCR to create the final RNA-seq library. The libraries were then quantified using a Qubit fluorometer and a TapeStation, pooled, and sequenced on an Illumina NovaSeq 6000 platform. Demultiplexing with bcl2fastq2 was performed to generate fastq files, with an average of 30 million reads per sample and three biological replicates per genotype. Reads were trimmed and aligned to the mouse reference genome (GRCm38/mm10) using STAR, and transcript abundance was estimated using featureCounts. Differentially expressed genes were identified with DESeq2 using a threshold of FDR < 0.05, and genes showing a |log2 fold change| > 1 were considered significantly changed. Gene Ontology and KEGG pathway enrichment analyses were performed using the clusterProfiler package to identify biological processes affected by *Cops5* deletion.

### 2.6 RT-qPCR

Total RNA was reverse-transcribed into cDNA using a reverse transcription kit. Quantitative PCR was performed using SYBR Green Master Mix on a real-time PCR system, and all primer used are listed in the **Supplemental Table S1**. Relative gene expression levels were calculated using the 2^^−ΔΔCt^ method, with 18S rRNA serving as an internal control. Each sample was analyzed in triplicate, and at least three biological replicates were included per group.

#### 2.6.1 Cell cycle analysis by flow cytometry

Testicular cell suspensions were prepared by enzymatic digestion with collagenase type IV and dispase, followed by mechanical disruption and filtration through a 40-μm cell strainer. Cells were fixed in ethanol, stained with propidium iodide, and analyzed on a flow cytometer. Single cells were gated on DNA content to quantify the relative proportions of cells in G1 (2C), S phase (2C–4C), and G2/M (4C) populations, using the DNA content of spermatozoa as the 1C reference.

### 2.7 Proteomics

Testes from 16-day-old control and cKO mice (n = 3 per group) were analyzed. Briefly, testicular proteins were extracted in a urea-containing lysis buffer, reduced, alkylated, and digested overnight with trypsin using a filter-aided sample preparation approach. Peptides were analyzed by liquid chromatography–tandem mass spectrometry (LC-MS/MS) using a high-resolution Orbitrap Fusion mass spectrometer at the Center for Metabolomics and Proteomics at the University of Minnesota. Raw data were processed using MaxQuant software, and label-free quantification was applied with a minimum peptide ratio count of 1. Proteins were identified with a false discovery rate of 1% at both the peptide and protein level. Differentially expressed proteins were identified using P < 0.05 and fold-change ≥ 2.0. Integrated analysis of proteomic and transcriptomic datasets was performed to identify concordant molecular changes.

### 2.8 Immunofluorescence

Testicular sections (5 μm) were subjected to antigen retrieval in citrate buffer, blocked with 5% normal goat serum for 1 h at room temperature, and incubated overnight at 4 °C with primary antibodies against transposon-related and PIWI proteins. After three washes, sections were incubated with fluorophore-conjugated secondary antibodies for 1 h at room temperature and then counterstained with DAPI. Sections were mounted in antifade mounting medium. Images were captured using a confocal microscope (Zeiss LSM 980) and processed with ZEN imaging software.

### 2.9 Small RNA sequencing

Total RNA was isolated from 16-day-old control and cKO mice and enriched for small RNAs (18–40 nt). Approximately 500 ng of total RNA was used for small RNA-seq library preparation by following the TriLink Cleantag small RNA-seq sample preparation guide (TriLink Biotechnologies, San Diego, CA). The CleanTag™ Small RNA Library Prep Kit uses a specialized, high-efficiency ligation strategy to minimize adapter-dimer formation for small RNA sequencing. The workflow involves 3’ adapter ligation, 5’ adapter ligation, reverse transcription, and PCR amplification. The PCR products were then purified to create the final small RNA-seq library. The small RNA-seq libraries were then quantified, pooled, and sequenced on an Illumina NovaSeq 6000 platform. Demultiplexing with bcl2fastq2 was then performed to generate fastq files, with an average of 25–30 million reads per sample and three biological replicates per genotype. After adapter removal, reads 18–40 nt in length were retained. Valid piRNAs were annotated as 24–32 nt small RNAs that map to the mouse genome and form RNA duplexes with PIWI proteins, as described previously. Reads were classified into piRNA, miRNA, mRNA, rRNA, tRNA, snRNA, snoRNA, and other categories, and the number of unique piRNA sequences was calculated after deduplication. Nucleotide composition at each position was analyzed across the first 15 nt of the valid piRNA population to assess the 1U bias.

### 2.10 Statistical analysis

All quantitative data are presented as mean ±SEM. Comparisons between two groups were conducted using two-tailed Student’s t-test, and comparisons involving more than two groups were analyzed by one-way analysis of variance followed by Tukey’s multiple-comparisons test. Each biological replicate represents an independent mouse, and n values are indicated in the corresponding figure legends. A p value < 0.05 was considered statistically significant.

## 3. Results

### 3.1 COPS5 is highly expressed in human germ cells and is reduced in patients with non-obstructive azoospermia

To evaluate the expression landscape of COPS5 in human testes, we analyzed publicly available single-cell RNA sequencing (scRNA-seq) datasets (GSE149512 and GSE182786) comprising 22 normal testicular samples and 7 samples from patients with non-obstructive azoospermia (NOA)^35,36^. COPS5 transcripts were detected in multiple testicular cell populations, including germ cells, Leydig cells, peritubular cells, and macrophages (**Figure 1**). Among these populations, germ cells exhibited the highest relative level of COPS5 expression in normal testes. Compared with normal controls, COPS5 expression in germ cells was significantly reduced by 56.4% in NOA samples, whereas the relative expression levels in Leydig, peritubular, and macrophage populations were higher in NOA samples (**Figure 1**). These results suggest that reduced COPS5 expression in germ cells is associated with impaired spermatogenesis in humans.

**Figure 1.**
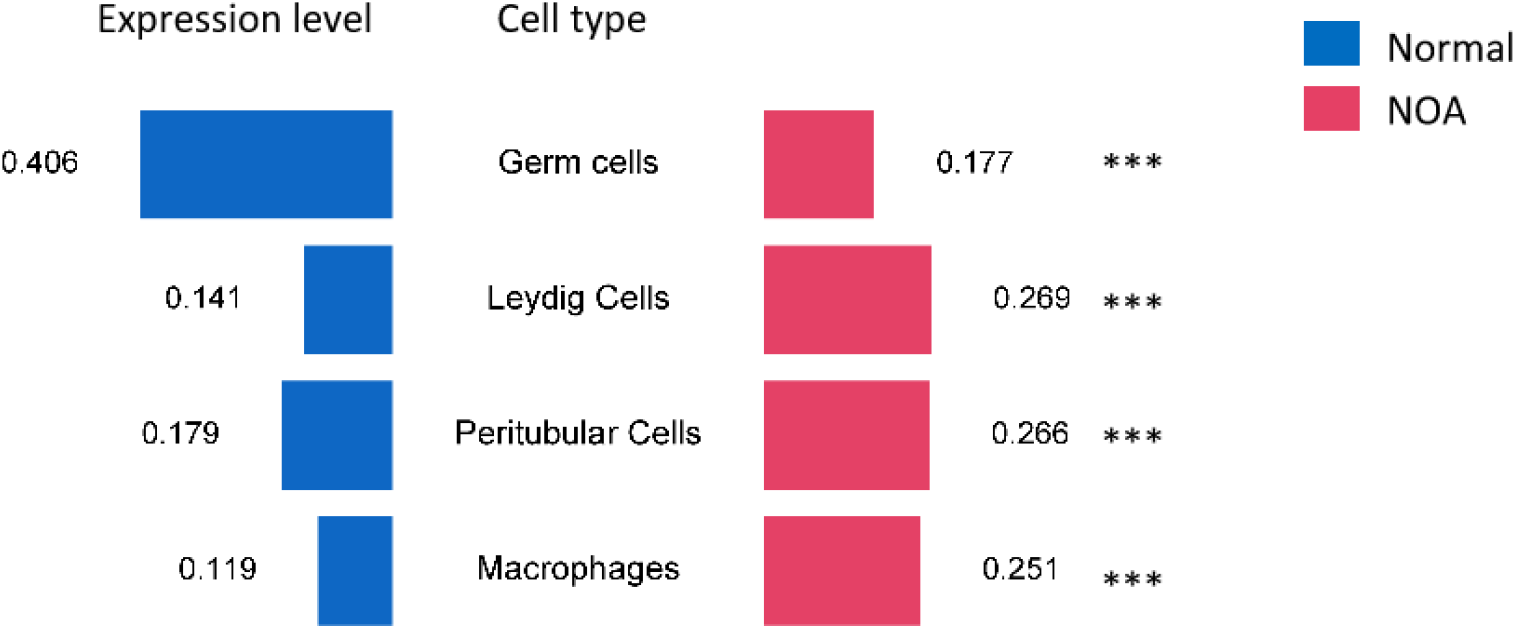
COPS5 expression in normal human testes and testes from patients with non-obstructive azoospermia (NOA) based on public single-cell RNA-seq datasets. Bar charts showing the relative COPS5 expression levels in the four indicated testicular cell types—germ cells, Leydig cells, peritubular cells, and macrophages—in normal controls (blue) and NOA patients (red). Data are presented as the mean values derived from the indicated public datasets (GSE149512 and GSE182786; n = 22 normal controls and 7 NOA patients). ***P < 0.001.

### 3.2 *Cops5* deficiency in germ cells causes progressive spermatogenic arrest after P16

COPS5 was detected in both the cytoplasmic and nuclear fractions of adult mouse testes by subcellular fractionation followed by Western blot analysis, with α-tubulin and Lamin B serving as the cytoplasmic and nuclear markers, respectively (**Figure 2A**). To further investigate the function of COPS5 during spermatogenesis, we used Stra8-iCre to generate germ cell-specific conditional gene knockout (cKO) mice of *Cops5* (*Cops5* cKO). To define the stage at which *Cops5* deletion first impairs spermatogenesis, testicular morphology was examined during the first wave of spermatogenesis (**Figure 2B**). No appreciable differences were observed between control and *Cops5* cKO mice from P8 to P16. By P20, however, the seminiferous epithelium of cKO mice began to disorganize. At P22, control testes contained abundant pachytene spermatocytes and nascent round spermatids, whereas cKO testes contained only scarce pachytene spermatocytes and were essentially devoid of round spermatids. By P28, control testes displayed the complete spectrum of germ cells, including spermatogonia, spermatocytes, round spermatids, and elongating spermatids, whereas elongating spermatids were absent in cKO testes. At P16, the organization of the seminiferous epithelium was not grossly altered in cKO testes, and the percentages of 2C and 4C cell populations, quantified on the basis of DNA content, were comparable between the two groups (**Figure 2C**). Collectively, these results indicate that *Cops5* deletion in germ cells leads to a progressive arrest of spermatogenesis after P16, coinciding with meiosis and the onset of spermiogenesis.

**Figure 2.**
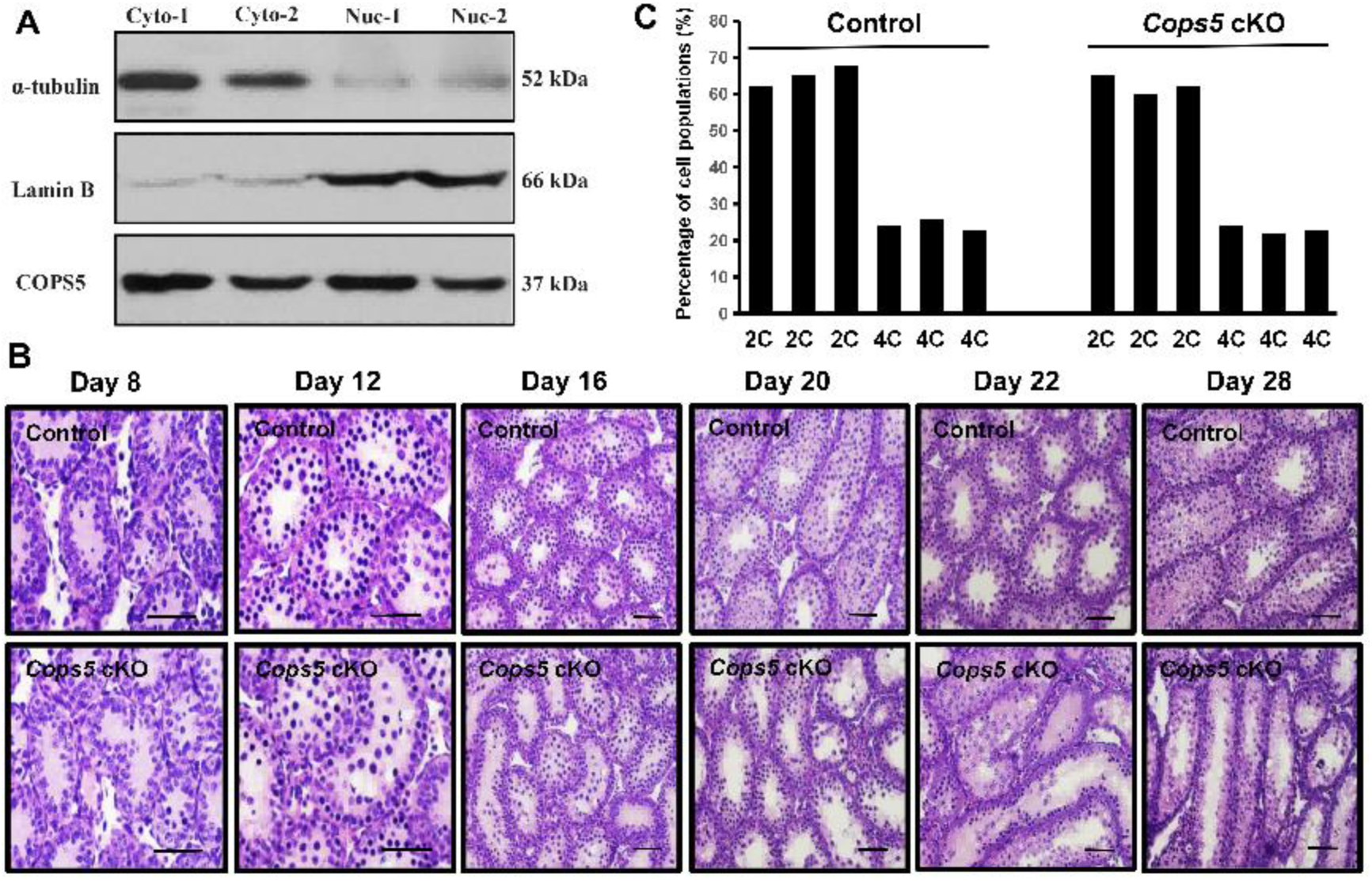
Onset of spermatogenic defects in germ cell-specific *Cops5* cKO mice. (A) Subcellular fractionation of adult testicular lysates followed by Western blot analysis showing that COPS5 is present in both the cytoplasmic (Cyto) and nuclear (Nuc) fractions; α-tubulin and Lamin B served as the cytoplasmic and nuclear markers, respectively. (B) Representative H&E-stained testicular sections from control and *Cops5* cKO mice at P8, P12, P16, P20, P22, and P28 during the first wave of spermatogenesis. Scale bars: 50 μm. (C) Quantification of the percentages of 2C and 4C cell populations, based on DNA content, in P16 control and *Cops5* cKO testes (n = 3 per group).

### 3.3 Loss of *Cops5* causes widespread transcriptomic dysregulation with prominent suppression of piRNA biogenesis and spermiogenesis-related programs

To define the molecular consequences of *Cops5* deletion, we performed RNA sequencing (RNA-seq) on P16 control and *Cops5* cKO testes. Differential expression analysis revealed extensive transcriptional dysregulation in cKO testes, with 9,912 differentially expressed genes (∣log2(FC)∣ > 0 and FDR < 0.05), among which 449 genes were upregulated and 1,641 genes were downregulated by more than twofold (∣ log2(FC)∣ > 1 and FDR < 0.05). Among them, key components of the piRNA pathway involved in germ cell development (*Piwil1/Miwi*, *Piwil2/Mili*, *Tdrd1*, *Tdrd5*, *Tdrd6*, *Tdrd9*, *Tdrkh*, *Mael*, *Ddx4, Mov10l1*, *Btbd18*, *Gasz*, *Rnf17*, *Stk31*, and *Pbk*), and genes related to cilium/flagellum function and intraflagellar transport (*Ift122, Ift140, and Wdr35*) were affected (**Figure 3A**). KEGG pathway enrichment analysis of the upregulated transcripts highlighted axon guidance, ovarian steroidogenesis, ErbB signaling, and PI3K-Akt signaling pathways (**Figure 3B**). Gene Ontology (GO) enrichment analysis of the downregulated genes identified terms in all three ontologies, including cilium movement, microtubule-based movement, cilium- or flagellum-dependent cell motility, axoneme assembly, cilium organization, and sperm motility **(Figure 3C**), consistent with the early spermatogenic defects observed in cKO testes. To validate the RNA-seq findings, representative differentially expressed genes were quantified by RT-qPCR using P16 testes. Consistent with the transcriptomic profiling, mRNA levels of piRNA pathway components, including *Piwil1, Piwil2, Btbd18, Tdrd5, Tdrd6, Mael, Gasz, Henmt1, Tdrkh, Spocd1,* and *Mov10l1*, as well as other spermatogenesis-related genes such as *Pgk2, Ift122, Ift140,* and *Fkbp6*, were significantly reduced in *Cops5* cKO testes (*P* < 0.05; **Figure 3D**).

**Figure 3.**
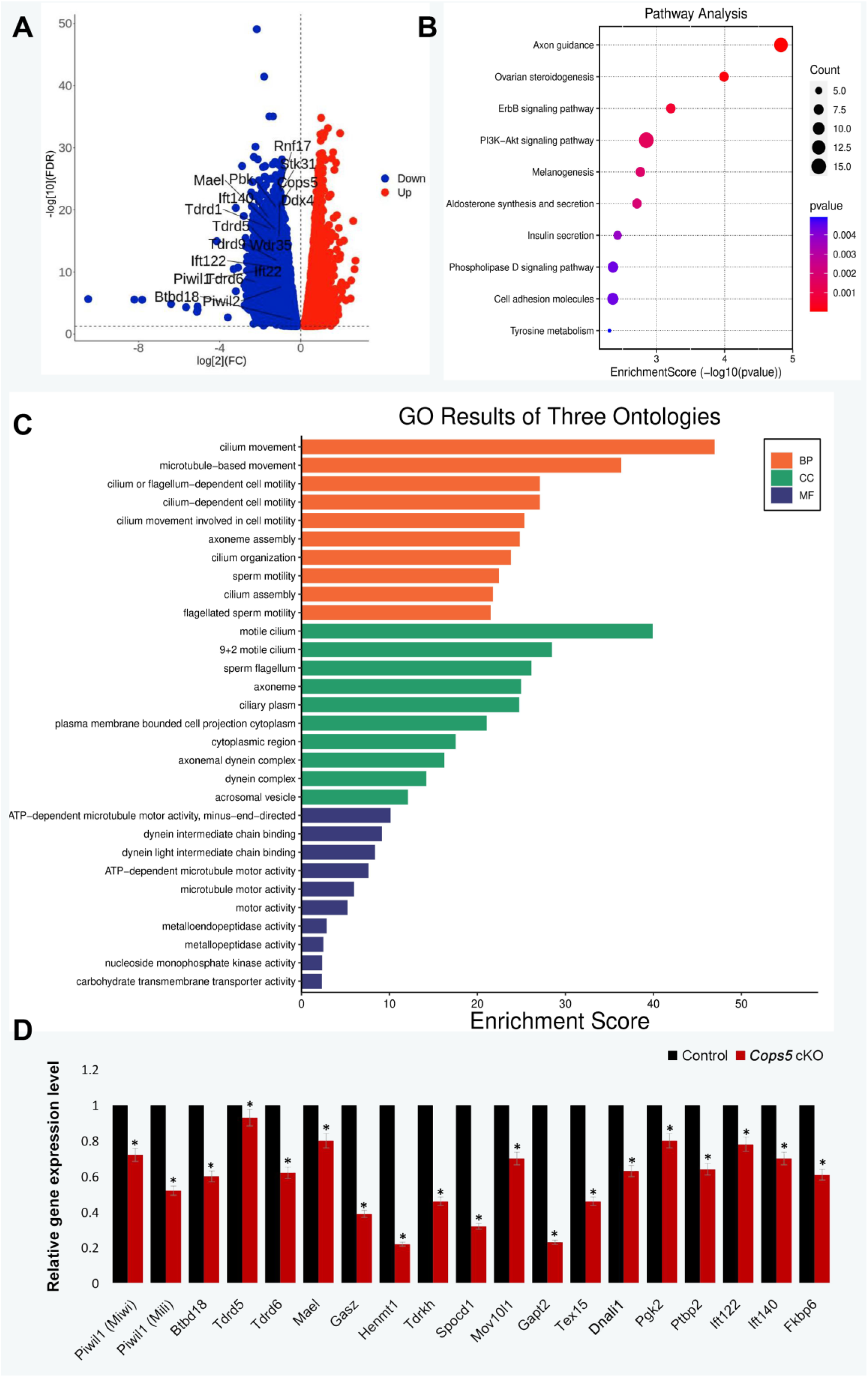
Transcriptomic profiling of P16 control and *Cops5* cKO testes. (A) Volcano plot of differentially expressed genes in P16 control and *Cops5* cKO testes; genes downregulated and upregulated in cKO testes are shown in blue and red, respectively (∣log2 (FC)∣ < 0 and FDR < 0.05); selected genes are labeled. (B) KEGG pathway enrichment analysis of the upregulated genes. (C) GO enrichment analysis (biological process, cellular component, and molecular function) of the downregulated genes. (D) RT-qPCR validation of selected downregulated genes, including piRNA pathway components (*Piwil1/Miwi, Piwil2/Mili, Btbd18, Tdrd5, Tdrd6, Mael, Gasz, Henmt1, Tdrkh, Spocd1,* and *Mov10l1*) and other spermatogenesis-related genes (*Tex15, Dnali1, Pgk2, Ptbp2, Ift122, Ift140*, and *Fkbp6*). Data are presented as mean ±SEM; *P < 0.05.

### 3.4 *Cops5* loss remodels the testicular proteome

Total testicular lysates from P16 control (n = 3) and *Cops5* cKO (n = 3) littermates were subjected to comparative proteomic profiling. Correlation analysis demonstrated distinct protein-expression patterns between control and cKO samples (**Figure 4A**). In total, 2,978 proteins exhibited altered abundance in cKO testes; applying thresholds of *P* < 0.05 and fold change ≥ 2.0, 65 proteins were significantly increased and 399 proteins decreased in cKO testes (**Figure 4B**). Protein–protein interaction (PPI) highlighted the piRNA metabolic process and DNA methylation, cilium assembly and intraciliary transport, RNA splicing and degradation, vesicle-mediated transport, and RNA transport (**Figure 4C**). To confirm these key findings, Western blot analysis was performed on P16 testicular lysates. Proteins closely associated with intraflagellar transport and flagellar assembly (IFT74 and IFT140, together with the IFT-associated regulator LZTFL1), the flagellar structural protein SPAG16, and proteins implicated in spermatid development and head–tail coupling (MEIG1, RIMBP3, and TCFL5) were markedly reduced in *Cops5* cKO testes (*P* < 0.05; Figure 4D, E), consistent with the proteomic profiling results.

**Figure 4.**
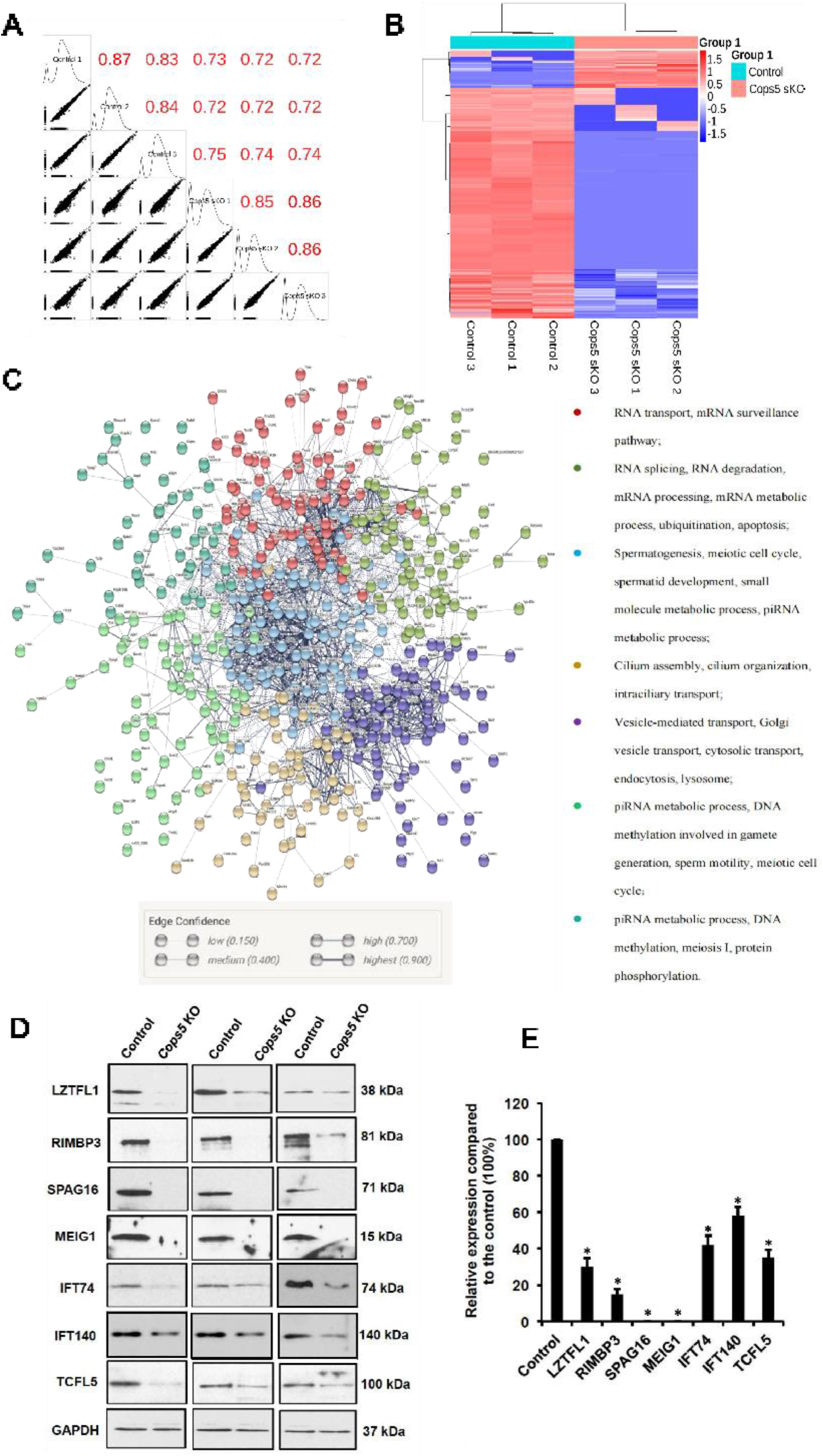
Comparative proteomic analysis of P16 control and *Cops5* cKO testes. (A) Correlation matrix showing the relationships among biological replicates for the control and *Cops5* cKO proteomes (n = 3 per group); Pearson correlation coefficients are indicated. (B) Heatmap summarizing the differentially abundant proteins between groups. (C) Protein–protein interaction (PPI) analysis of the differentially expressed proteins, color-coded by functional category as indicated. (D, E) Western blot analysis (D) and quantification (E) of proteins closely related to intraflagellar transport and spermatid development (LZTFL1, RIMBP3, SPAG16, MEIG1, IFT74, IFT140, and TCFL5); GAPDH served as the loading control. Data are presented as mean ± SEM (n = 3); *P < 0.05.

### 3.5 Integrated transcriptomic and proteomic analyses identify the piRNA pathway as a core COPS5-regulated module

To identify transcripts and proteins that were consistently affected, RNA-seq and proteomic datasets from P16 testes were combined revealing 59 gene protein pairs with concurrently reduced in cKO testes. In contrast, only two gene protein pairs were concordantly upregulated (**Figure 5A**). GO biological process enrichment analysis of the 59 concordantly downregulated genes showed significant enrichment in terms related to the piRNA metabolic process, the meiotic cell cycle, DNA methylation involved in gamete generation, germ cell development, spermatid development and differentiation, and intraciliary transport (**Figure 5B**). Network visualization further highlighted representative factors within these modules, including the PIWI protein PIWIL1, multiple Tudor-domain proteins (TDRD1, TDRD5, and TDRD6), the RNA helicases DDX4 and DDX25, the piRNA biogenesis factor MAEL, the meiosis-associated protein MEIG1, and the intraflagellar transport protein IFT140 (**Figure 5C**). In the RNA-seq dataset, all of these genes were reduced by more than two-fold in cKO testes (**Figure 5D**). Given the consistent suppression of piRNA-associated transcripts and proteins, the expression of key piRNA pathway proteins was further examined by Western blotting. At P16, TDRD1, TDRD5, DDX4, MILI (PIWIL2), and MYBL1 were all significantly reduced in cKO testes (*P* < 0.05; **Figure 5E, F**). Consistently, the immunofluorescence signals of the PIWI proteins PIWIL1/MIWI and PIWIL2/MILI were markedly reduced in cKO testes (**Figure 6**; **Supplemental Figures S1 and S2**), This level of consistently, strongly supports the role of COPS5 in maintaining the piRNA pathway at the protein level.

**Figure 5.**
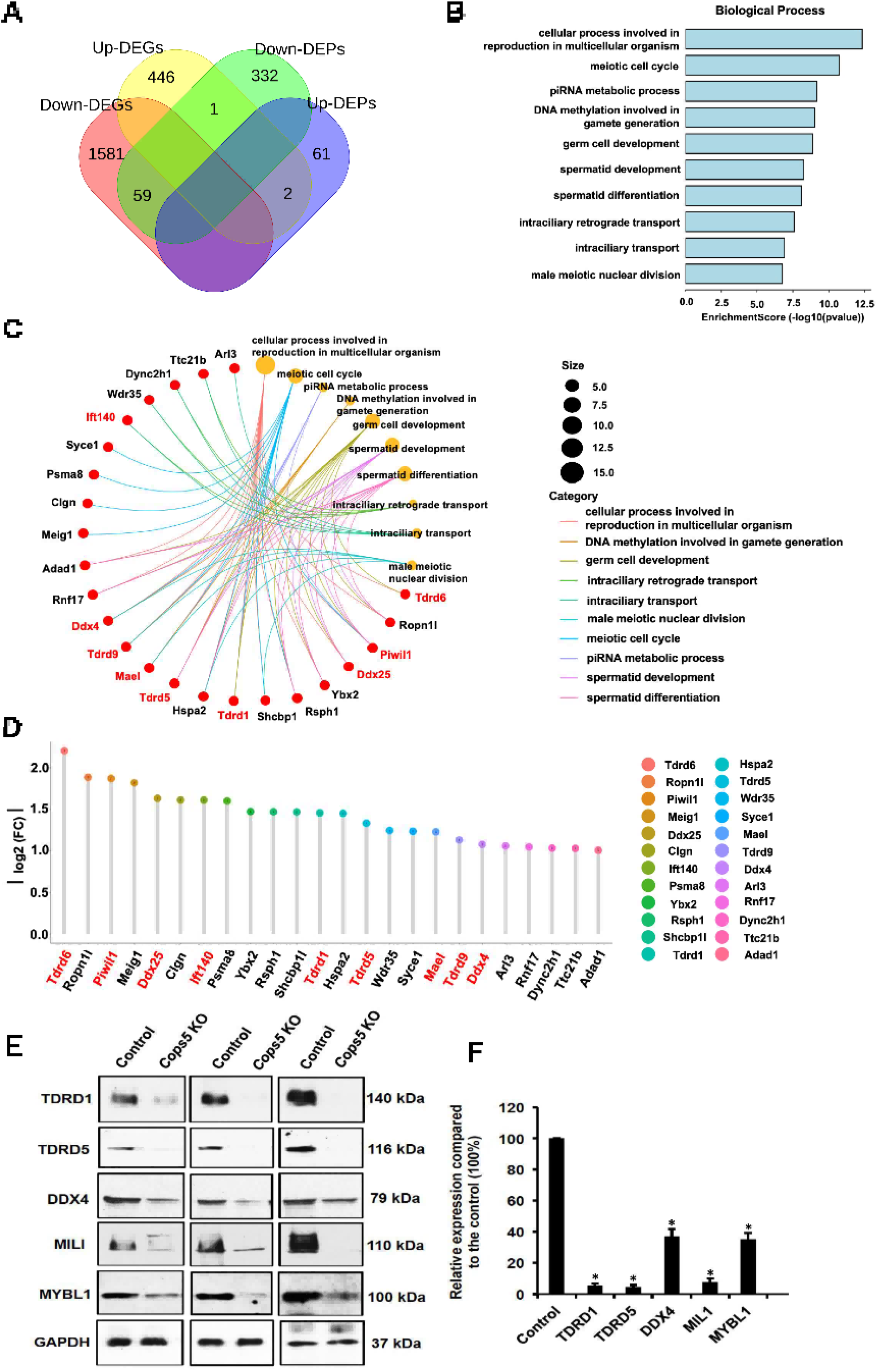
Integrated transcriptomic and proteomic analyses identify core COPS5-regulated factors. (A) Venn diagram showing the overlap between downregulated genes (Down-DEGs, 1581), upregulated genes (Up-DEGs, 446), downregulated proteins (Down-DEPs, 332), and upregulated proteins (Up-DEPs, 61) in P16 *Cops5* cKO testes; 59 genes were concordantly downregulated at both the mRNA and protein levels, and 2 genes were concordantly upregulated. (B) GO biological process enrichment analysis of the 59 concordantly downregulated genes. (C) cnet plot illustrating the connections between enriched terms and the associated genes. (D) Lollipop chart summarizing the |log2 (FC)| values of the concordantly downregulated genes. (E, F) Western blot analysis (E) and quantification (F) of piRNA pathway proteins (TDRD1, TDRD5, DDX4, MILI, and MYBL1) in P16 control and *Cops5* cKO testes; GAPDH served as the loading control. Data are presented as mean ±SEM (n = 3); *P < 0.05.

**Figure 6.**
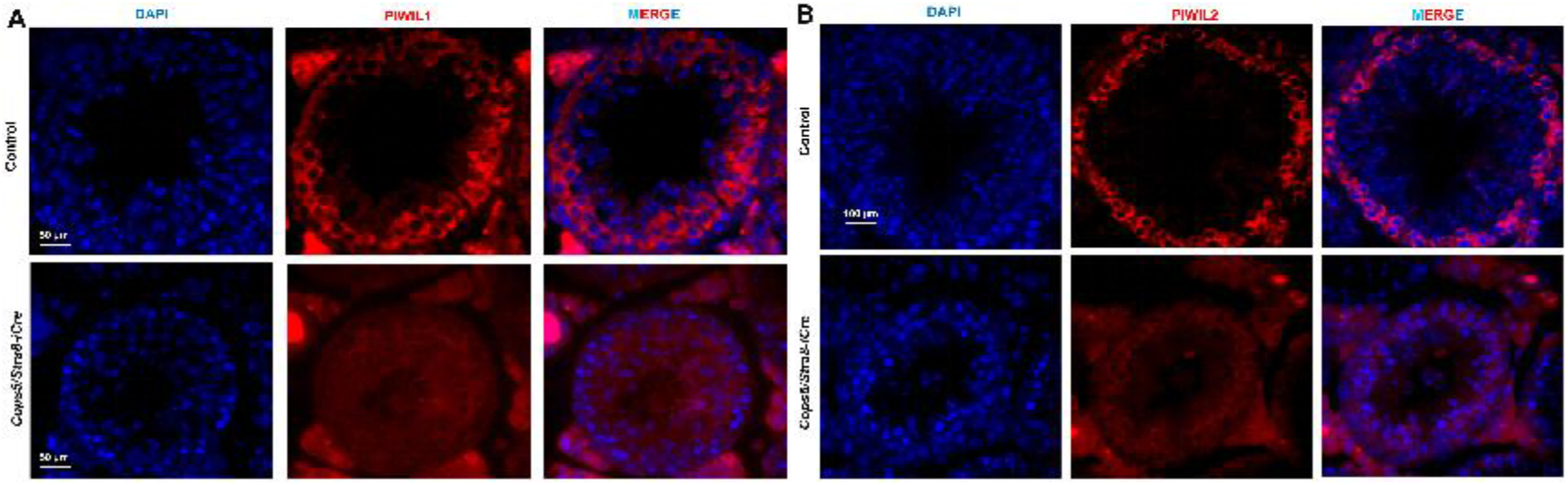
Conditional knockout of *Cops5* decreases PIWIL1 (MIWI) and PIWIL2 (MILI) expression in the testis. (A, B) Representative immunofluorescence images showing the expression of PIWIL1 (A) and PIWIL2 (B) in testicular sections from control and *Cops5/Stra8-iCre* (cKO) mice; nuclei were stained with DAPI (blue). Scale bars: 50 μm (A) and 100 μm (B).

### 3.6 Small RNA sequencing reveals impaired pachytene piRNA biogenesis in *Cops5* cKO testes

The results of the combined omics analysis indicated that the deletion of *Cops5* disrupted the production of piRNAs. To confirm the above, small RNA sequencing of P16 control and cKO testes (n = 3 per group) was carried out. Length-resolved classification of small RNAs revealed a pronounced reduction in the 24–32-nt piRNA population in cKO testes, accompanied by a marked relative increase in approximately 22-nt miRNAs (**Figure 7a–f**). Consistent with this shift, piRNAs accounted for approximately 56% of total small RNAs in control testes but only 18–30% in cKO testes (**Figure 7g–l**). *Cops5* deletion reduced not only the overall abundance of piRNAs but also the number of unique piRNA sequences across the entire 24–32-nt range, indicating a marked reduction in the diversity of the piRNA repertoire (**Figure 7m–s**). Despite these quantitative defects, the nucleotide composition of the remaining piRNAs was broadly preserved, including the characteristic 1U bias (**Figure 7t, u**). Together, these findings demonstrate that COPS5 is required for both the abundance and the diversity of pachytene piRNAs, supporting an essential role for *COPS5* in pachytene piRNA biogenesis.

**Figure 7.**
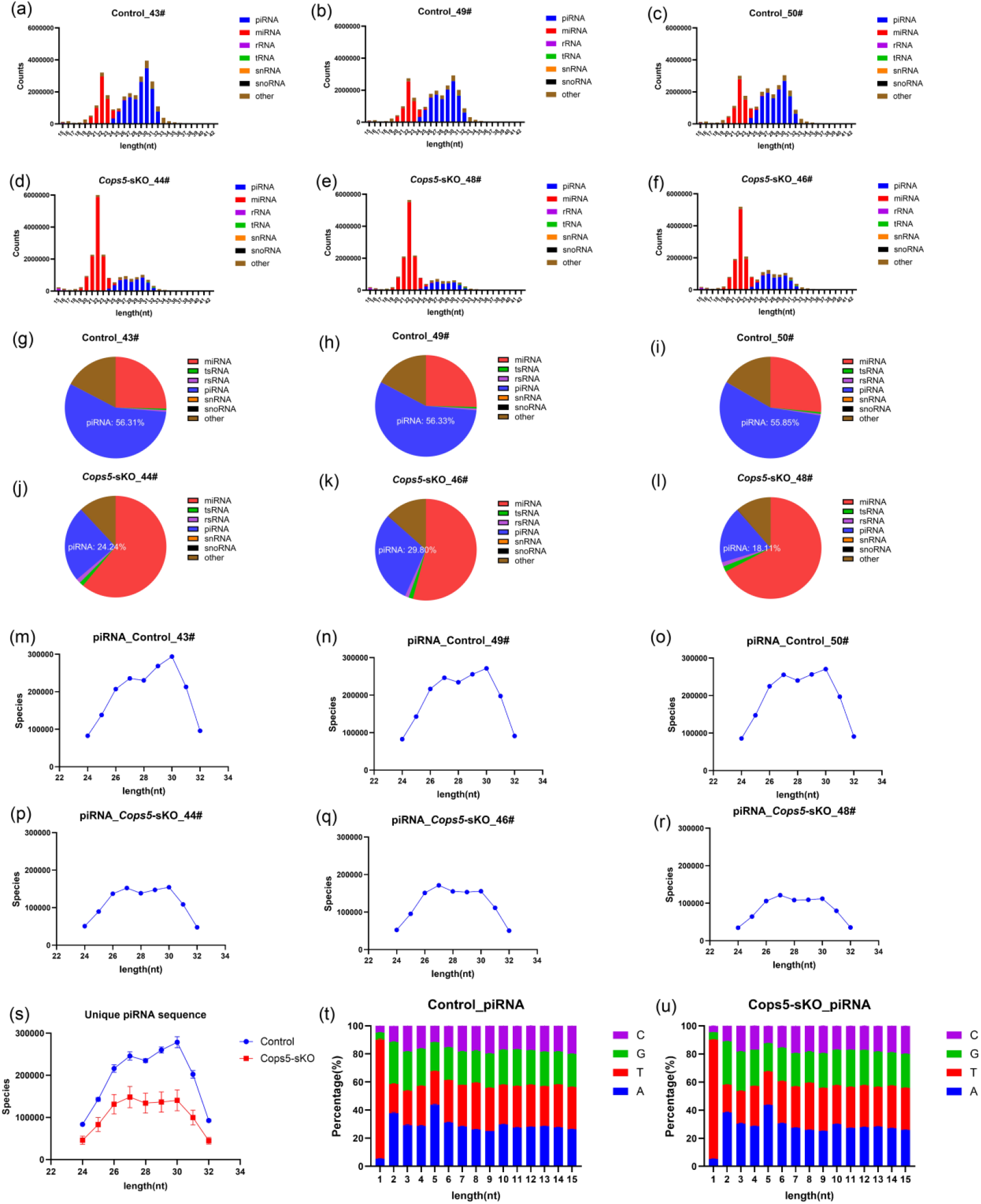
Loss of COPS5 impairs pachytene piRNA abundance and diversity in the testis. (a–f) Length-resolved classification of small RNAs (piRNA, miRNA, mRNA, rRNA, tRNA, snRNA, snoRNA, and other) from small RNA sequencing of individual P16 control (a–c) and *Cops5* cKO (d–f) testes. (g–l) Relative proportions of small RNA classes in individual control (g–i) and *Cops5* cKO (j–l) testes; the percentage of piRNAs is indicated. (m–r) Numbers of unique piRNA sequences across the 24–32-nt range in individual control (m–o) and *Cops5* cKO (p–r) testes. (s) Comparison of unique piRNA sequences between control and *Cops5* cKO testes (mean ± SEM). (t, u) Nucleotide composition at positions 1–15 of the remaining piRNAs in control (t) and *Cops5* cKO (u) testes, showing retention of the 1U bias. n = 3 mice per group.

### 3.7 *Cops5* deficiency causes early activation of transposable elements

piRNAs safeguard germline genome integrity, in part by repressing transposable elements (TEs). Although the cellular composition of P16 testes was comparable between control and cKO testes (**Figure 2C**), we examined the earlier time point of P14 to exclude potential confounding effects of altered cell composition. Western blot analysis of P14 testes showed that multiple piRNA pathway proteins, including TDRD1, TDRD5, TDRD7, MAEL, MYBL1, and MILI, were already significantly reduced in *Cops5* cKO testes, whereas the retrotransposon IAP protein was significantly elevated (**Supplemental Figure S3**), indicating an early, *Cops5*-dependent impairment of the piRNA pathway that precedes overt spermatogenic failure. At P16, immunofluorescence staining showed markedly enhanced LINE1 signals in cKO seminiferous tubules (**Figure 8A**, **Supplemental Figure S4**), and Western blot analysis confirmed significant increases in LINE1 (**Figure 8B**) and IAP (**Figure 8C**) protein levels. Together, these data demonstrate that *Cops5* loss leads to activation of TE expression, consistent with compromised piRNA-mediated transposon silencing.

**Figure 8.**
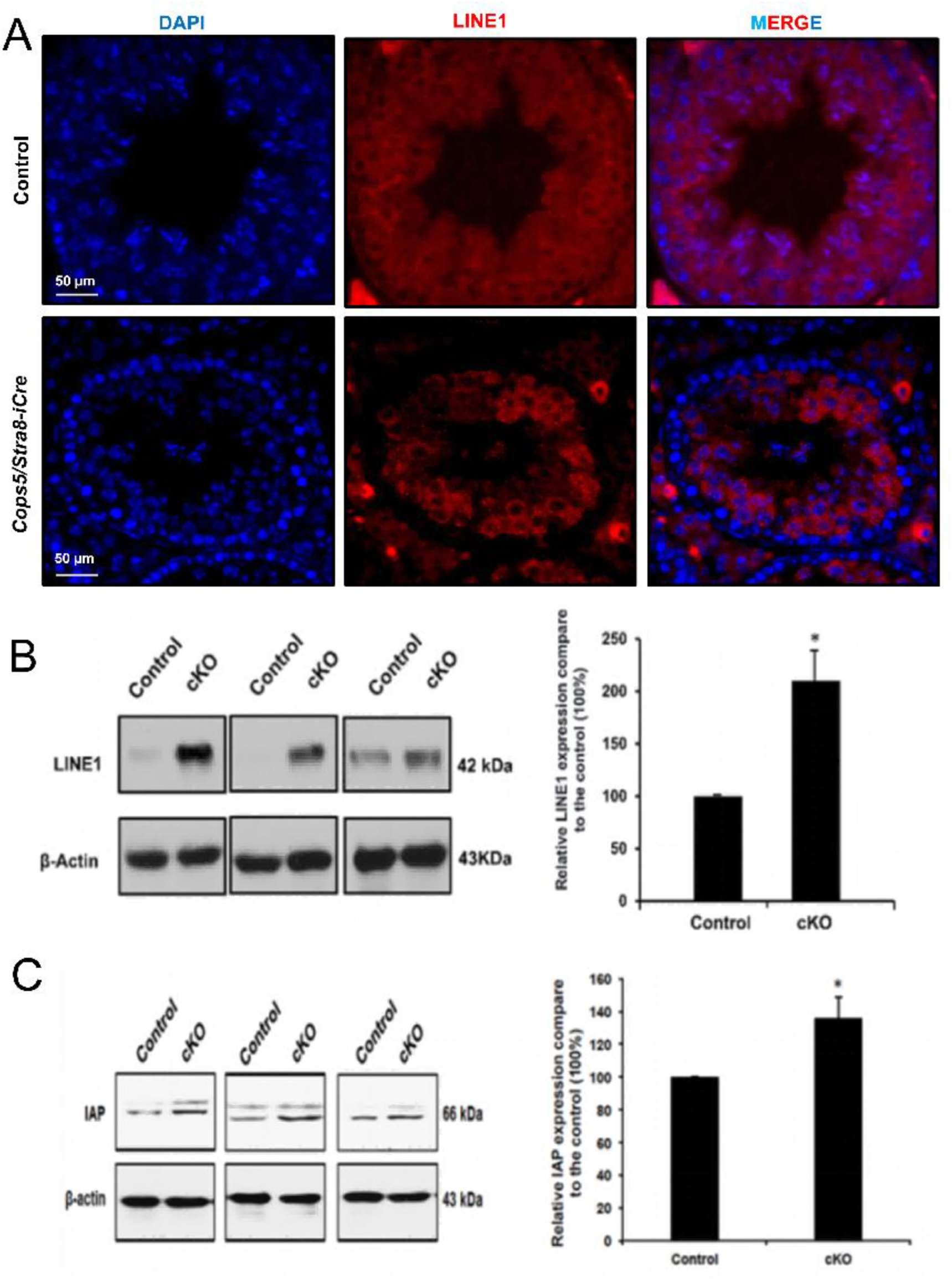
Conditional knockout of Cops5 increases LINE1 and IAP retrotransposon expression in mouse testes. (A) Representative immunofluorescence images showing markedly enhanced LINE1 signals (red) in seminiferous tubules of *Cops5/Stra8-iCre* (cKO) testes compared with controls; nuclei were stained with DAPI (blue). Scale bar, 50 μm. (B, C) Western blot analysis and quantification of LINE1 (B, 42 kDa) and IAP (C, 66 kDa) in P16 control and *Cops5* cKO testes, with β-actin as the loading control. Data are presented as mean ± SEM (n = 3 mice per group); *P < 0.05.

## 4. Discussion

While the cellular compositions at day 16 after birth of the COPS5-control and cKO mice are similar, the outcomes differ. To address the basis, an integrated multi-omics strategy was employed. This began to define the molecular mechanism by which COPS5 informs male germ cell development. RNA-seq, proteomics and small RNA-seq identified the piRNA biogenesis pathway as a major COPS5-dependent module. Small RNA sequencing showed that both the abundance and the sequence diversity of pachytene piRNAs were markedly reduced in *Cops5* cKO testes. Consistent with impaired piRNA production, levels of LINE1 and IAP retrotransposons increased. Interestingly, COPS5 expression was reduced in the germ cells of patients with non-obstructive azoospermia (NOA). These findings emphasize that COPS5 is required for pachytene piRNA biogenesis and transposon silencing during male germ cell development.

The piRNA pathway is an evolutionarily conserved defense system that protects germline genome integrity by silencing transposable elements (TEs) ^13–15^. Mouse piRNAs are produced in two temporally distinct waves: fetal piRNAs that establish DNA methylation-based silencing ^16–19^ and postnatal pachytene piRNAs that act mainly at the post-transcriptional level ^20^. Loss of function of individual piRNA pathway components in mice leads to TE activation, meiotic defects, and male infertility ^21–32^.

Our data are consistent with this framework. In *Cops5* cKO testes, many piRNA pathway genes were coordinately reduced at both the mRNA and protein levels. Because transcripts and proteins changed in parallel, the piRNA defect in *Cops5* cKO testes is more likely to reflect a broad impairment of the gene expression program that supports piRNA biogenesis than the loss of a single factor. The phenotype of *Cops5* cKO mice, which includes a defect in meiotic progression and the loss of round and elongating spermatids, closely resembles that of mice lacking *Mili, Miwi, Mael, Tdrd1, Tdrd5, Tdrd9, Mov10l1, Tdrkh, Pnldc1, Henmt1,* or *Tex15* ^21–32^, supporting the view that COPS5 acts within or upstream of the piRNA pathway.

Small RNA sequencing provided direct evidence for a defect in piRNA production. The proportion of piRNAs decreased from approximately 56% of total small RNAs in control testes to 18-30% in *Cops5* cKO testes, and the number of unique piRNA sequences across the 24-32 nt range was markedly reduced, indicating that both the amount and the diversity of the pachytene piRNA pool were affected. In contrast, the characteristic 1U bias of the remaining piRNAs was largely preserved. A similar dissociation between piRNA abundance and 1U bias has been reported in mice lacking *Tdrkh, Pnldc1,* or *Henmt1*, in which the final steps of piRNA maturation are impaired while earlier steps of piRNA production remain at least partly active ^29–31^. Our results therefore suggest that COPS5 is required for the efficient production or stabilization of mature pachytene piRNAs rather than for the machinery that establishes 1U identity. Because *Cops5* cKO testes contain fewer pachytene spermatocytes at P16 and later, part of the reduction in piRNA abundance may also reflect a smaller number of piRNA-producing cells; however, the cellular composition of P16 testes was comparable between genotypes, and TE activation was already detectable at P14, which suggests that the piRNA defect is cell-intrinsic rather than a simple consequence of cell loss.

Consistent with impaired piRNA production, the expression of two representative TEs, LINE1 and IAP, was increased in *Cops5* cKO testes. Both elements are physiological targets of the piRNA pathway in the male germline. In fetal germ cells, piRNA-guided DNA methylation of LINE1 and IAP regulatory regions by MILI and MIWI2 is required for their silencing ^16,17^. In adult germ cells, piRNAs bound to MILI and MIWI also suppress LINE1 at the post-transcriptional level, and this function is required for meiotic progression ^20,37,38^. A widely discussed mechanism that links TE activation to meiotic arrest involves the LINE1-encoded proteins ORF1p and ORF2p. ORF1p packages LINE1 RNA into ribonucleoprotein particles, whereas ORF2p provides endonuclease and reverse transcriptase activities ^39^ and can generate DNA double-strand breaks ^40^. Such damage may activate DNA damage checkpoints and interfere with meiotic progression, providing a plausible link between piRNA loss, TE activation, and the spermatogenic arrest observed in *Cops5* cKO testes. However, whether TE activation directly causes the meiotic arrest in this model remains to be established.

Two coexisting mechanisms may account for the requirement for COPS5 in spermatogenesis. Mechanistically, COPS5 acts through the piRNA pathway to maintain TE silencing. COPS5 contains a JAMM metalloenzyme motif that removes the ubiquitin-like protein NEDD8 from cullin proteins, thereby regulating the activity of cullin-RING ubiquitin ligases ^8^. Through this activity, COPS5 influences the stability of many proteins. Our observation that piRNA pathway mRNAs and proteins were both reduced suggests that COPS5 regulates piRNA pathway gene expression broadly rather than acting on a single downstream factor. Consistent with this possibility, COPS5 was originally identified as JAB1, a coactivator that binds the activation domain of c-Jun and JunD and increases the specificity of AP-1-dependent transcription, and it interacts with other transcription factors, including p53, NF-κB, HIF-1α, SMAD4, and SMAD5^9,10^. In addition, MYBL1 (A-MYB), a transcription factor that drives the expression of pachytene piRNA precursors and of many piRNA biogenesis genes^41^, was markedly reduced in *Cops5* cKO testes (Figure 5E, F). These observations raise the possibility that COPS5 supports the activity of transcription factors that control the expression of piRNA pathway genes, and that the loss of this support contributes to the coordinated downregulation of piRNA biogenesis at both the mRNA and protein levels. Because the CSN is also a central regulator of protein ubiquitination, the effect of COPS5 loss may in addition involve changes in protein stability. Which of these levels predominates, and how COPS5 recognizes its substrates in germ cells, remain to be determined.

COPS5 may also regulate spermatogenesis-related genes independently of the piRNA pathway. Besides piRNA pathway genes, several genes involved in cilium and flagellum assembly and in intraflagellar transport (IFT), including *Ift122, Ift140, Wdr35, Spag16,* and *Meig1*, were reduced in *Cops5* cKO testes, and Gene Ontology terms such as cilium movement, axoneme assembly, and sperm motility were enriched among the downregulated genes. These genes are not classical components of the piRNA pathway, and their reduced expression is difficult to explain solely by TE activation. However, they are consistent with the previously reported interaction between COPS5 and IFT20 and with the acrosome and flagellum defects in *Cops5* mutant mice ^11^, and with the essential role of MEIG1 in manchette formation and sperm head development ^42^. It is therefore possible that COPS5 controls two partially separable processes: TE silencing through the piRNA pathway, and the expression of genes required for sperm morphogenesis. We emphasize that this second mechanism remains a hypothesis. In the present study we did not experimentally separate the direct from the indirect effects of COPS5 loss, and the downregulation of these genes may also be a secondary consequence of the arrest of germ cell differentiation.

Based on these results, we propose a working model for the function of COPS5 in spermatogenesis (**Figure 9**). In wild-type germ cells, COPS5 maintains the expression of piRNA biogenesis genes such as *Tdrd1* and *Tdrd5*, which ensures piRNA production, transposon methylation, and silencing of LINE1 and IAP. COPS5 also maintains the expression of spermatogenesis-related genes such as *Meig1* and *Pacrg*, which are required for meiosis and sperm development. In *Cops5* cKO germ cells, loss of COPS5 inhibits piRNA biogenesis and decreases transposon methylation, leading to de-repression of LINE1 and IAP; at the same time, the expression of spermatogenesis-related genes becomes aberrant. Together, these defects block meiosis, impair spermatogenesis, and finally cause male infertility.

**Figure 9.**
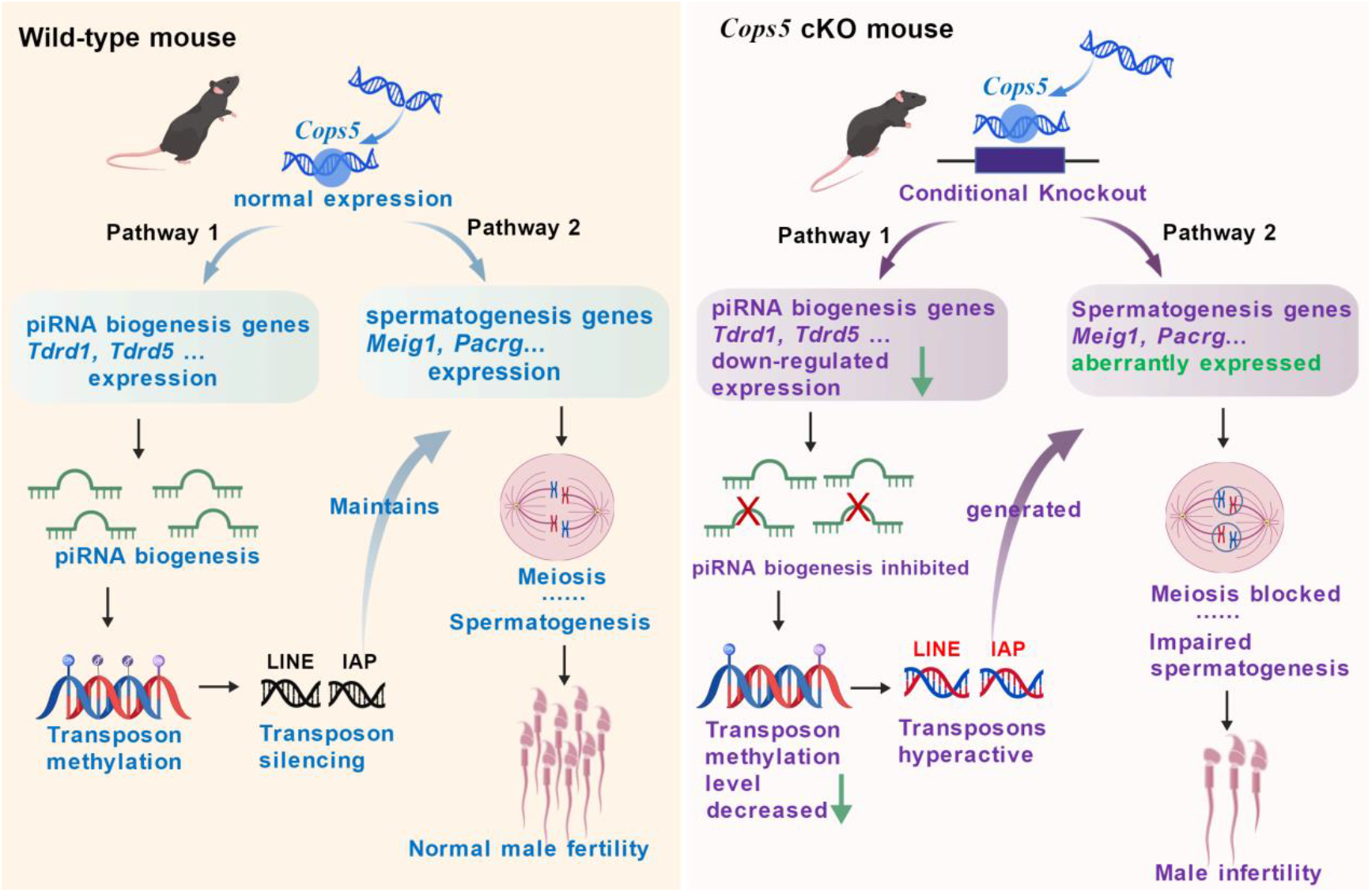
Proposed mechanism of COPS5 function in spermatogenesis. Left, in wild-type germ cells, COPS5 sustains the expression of piRNA biogenesis genes (*Tdrd1, Tdrd5*) and of spermatogenesis-related genes (*Meig1, Pacrg*), thereby ensuring piRNA production, transposon methylation, silencing of LINE1 and IAP, and normal meiosis and sperm development. Right, in Cops5 cKO germ cells, loss of COPS5 inhibits piRNA biogenesis and reduces transposon methylation, leading to de-repression of LINE1 and IAP and to abnormal expression of spermatogenesis-related genes; meiosis is blocked, spermatogenesis is impaired, and male infertility results.

Several limitations of this study should be acknowledged. First, our analysis was restricted to a limited number of developmental time points, and a detailed time course from P14 to P18 will be needed to determine whether piRNA loss and TE activation precede the arrest of meiosis or occur in parallel with it. Second, although the parallel changes in mRNA and protein suggest that COPS5 influences the piRNA pathway gene expression program, we did not identify the direct targets of COPS5, and the molecular link between COPS5 and piRNA biogenesis remains to be defined. Third, we did not examine DNA methylation of TE regulatory regions in *Cops5* cKO testes; such analysis, together with the quantification of Line1-Orf1 and Line1-Orf2 transcripts and of DNA damage markers such as γ-H2AX, would help to define how TE de-repression affects germ cell development.

Taken together, our results identify COPS5 as an essential regulator of the piRNA pathway and of transposon silencing during mouse spermatogenesis. We propose that loss of COPS5 impairs the production of pachytene piRNAs, leading to activation of LINE1 and IAP and to a progressive arrest of spermatogenesis. In addition, COPS5 may modify the expression of genes required for sperm morphogenesis. These findings extend the known functions of COPS5 in the male reproductive system and suggest that COPS5 dysfunction may contribute to human male infertility.

## 5. Conclusion

In this study, germ cell-specific deletion of *Cops5* in mice caused a progressive arrest of spermatogenesis, a marked reduction in the abundance and sequence diversity of pachytene piRNAs, and increased expression of the retrotransposons LINE1 and IAP. COPS5 expression was reduced in the germ cells of patients with non-obstructive azoospermia, suggesting that reduced COPS5 levels may contribute to human spermatogenic failure and may serve as a potential target for its diagnosis and treatment.

## Data Availability

The data is available from the corresponding author upon reasonable request.

## Acknowledgement

This research was supported by Wayne State University Start-up fund, NIH grants HD105944 (ZZ), HD107579 (ZZ), HD114311 (ZZ), and NIGMS R35GM124896 (YC). RNA-seq and small RNA-seq data were generated in the Genome Sequencing Facility which is supported by UT Health San Antonio, NIH-NCI P30 CA054174 (Cancer Center at UT Health San Antonio) and NIH Shared Instrument grant S10OD030311 (S10 grant to NovaSeq 6000 System), and CPRIT Core Facility Award (RP220662). We also thank Phillip D. Zamore at University of Massachusetts Medical School for discussion and suggestions on this project; Shinichiro Chuma at Kyoto University for providing TDRD1 and 7 antibodies; Alex Bortvin at Carnegie Institution for Science for providing MAEL antibody; Rolf Jessberger at Dresden University of Technology for providing TDRD6 antibody; Sandra L Martin at University of Colorado School of Medicine for providing LINE-1/IAP antibodies.

