## Supplementary Files for "COPS5 maintains spermatogenesis gene expression and piRNA-mediated transposon silencing"

USA

**Supplementary Table S1.** Primer sequences for RT-qPCR

| Gene | Forward primer (5′–3′) | Reverse primer (5′–3′) |
|---|---|---|
| *Piwil1* | CTCAAGTCAGTCGGGAGAGG | CAGGCCACTGCTGTCATTGA |
| *Piwil2* | ACAAATTGTTCGTTGCATCA | TTCAGCGGAATATCCACTCC |
| *Btbd18* | GGACTCTTCAGTGCTACCTGAC | AGGCTATATGAGACCCGTACAAC |
| *Tdrd5* | GCCCGATGTGGTTAGTGTCT | TATGCTGGGTCTTCCCTTTG |
| *Tdrd6* | ACCGTCACCCGATTAGACAG | AATGTTTCCCTCCCCCATTC |
| *Mael* | TTCCACAGGAGATTCGATTC | TCCATACGCTTCAAACACCA |
| *Gasz* | GGTGCTAATGCCAGCTTTGA | GCAGTATTTGGGTCCGCATT |
| *Henmt1** | GGATCCAGGAGTGGTCCAG | CTCCTCACGACTCAAGTGCT |
| *Tdrkh* | GAGTGCTGCTTATCAGTGGGT | CACCGCCTCTCCCTATGATT |
| *Spocd1* | ACCAACCTCCGCTACAAGAA | ATGTCCAGGCCCTTTCTCTC |
| *Mov10l1* | TTGGCTCGGTAGAGGAGTTC | TCTGACAAGCACATGAGGGT |
| *Gpat2* | AGTTGCTGAGGAGACTCCAG | CGGCAGAGGAAGAGGAAGAA |
| *Tex15* | TCCAGGTACAGAGTGCACAG | GGAGATCCTGTAGTGCTTGGA |
| *Dnali1* | GCTATGGATCCAGCAGGTGT | CGGATCAGTTCGTCAAAACA |
| *Ptbp2* | GCTTCCTCTGTTGCCAGTTC | TGCAGAAGACAAAAGCATGG |
| *Pgk2* | TTCATTGTGGAGGAAGAAGG | GGAAGGTGTAAGCCATTCCA |
| *Ift122* | CGAAGGAACTGCTTTTCCAG | GGTACATTGGAGCCGACTGT |
| *Ift140* | TTGAAAAGCTGTGGAGCTTT | CTTGGTGGCCAGGTGATAGT |
| *Fkbp6* | CACCGTCCTGTTTGAGATCG | CTTTGGCATCACAGAAGCGA |
| 18S rRNA | GGGAGCCTGAGAAACGGC | GGGTCGGGAGTGGGTAATTT |

**Supplemental figure**

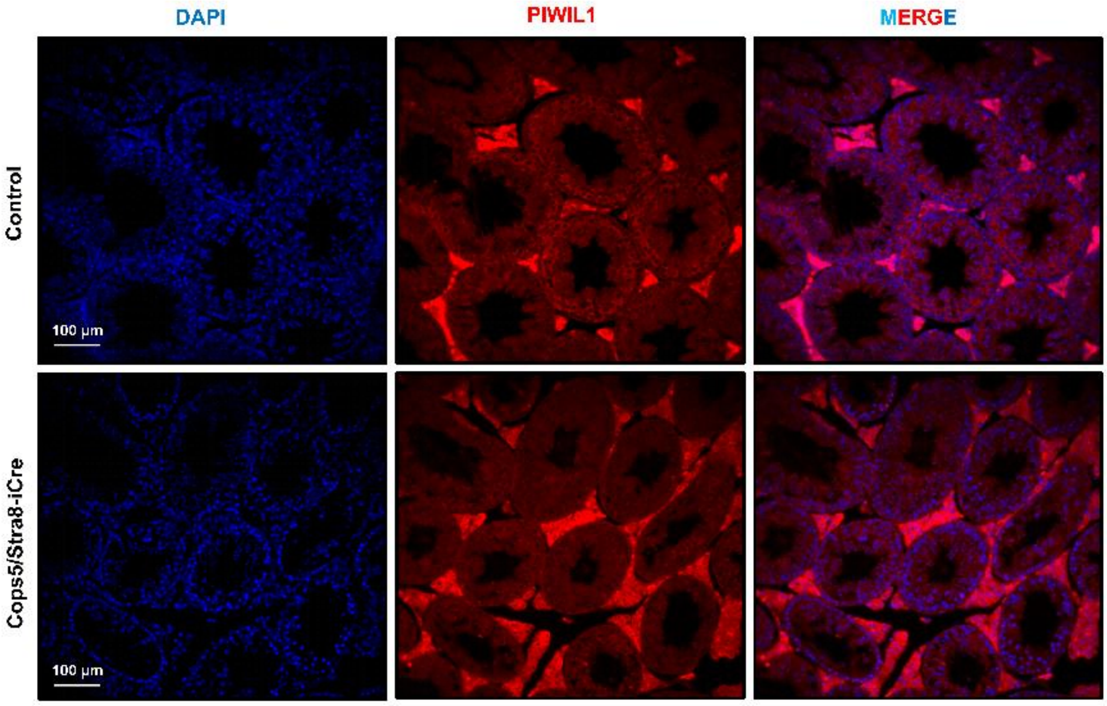


**Figure S1. Low-magnification immunofluorescence analysis of PIWIL1 in control and *Cops5* cKO testes.** Representative low-magnification images showing PIWIL1 immunoreactivity (red) in testis sections from control and *Cops5/Stra8-iCre* (cKO) mice; nuclei were counterstained with DAPI (blue). Scale bar, 100 μm.

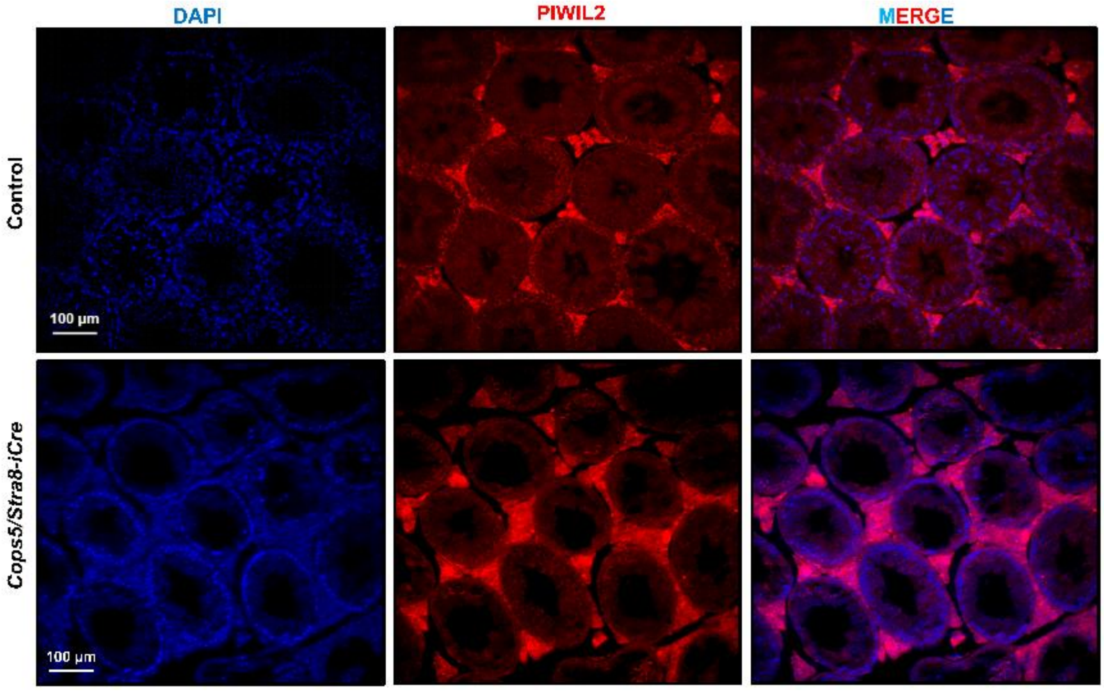

**Figure S2. Low-magnification immunofluorescence analysis of PIWIL2 in control and *Cops5* cKO testes.** Representative low-magnification images showing PIWIL2 immunoreactivity (red) in testis sections from control and *Cops5/Stra8-iCre* (cKO) mice; nuclei were counterstained with DAPI (blue). Scale bar, 100 µm.

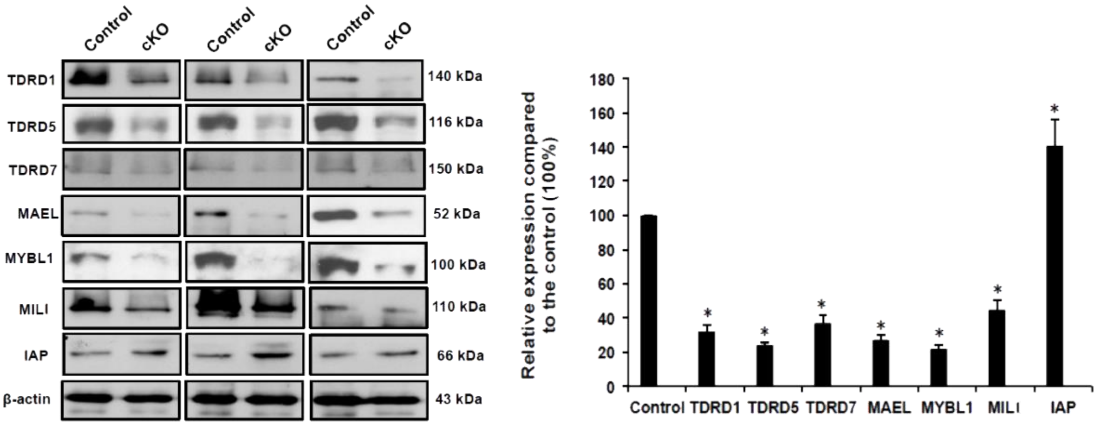


**Figure S3. Expression of piRNA pathway factors and the IAP retrotransposon protein in control and Cops5 cKO mice on P14.** Representative Western blots of the indicated proteins (TDRD1, TDRD5, TDRD7, MAEL, MYBL1, MILI, and IAP) in testis lysates from control and *Cops5* cKO mice; β-actin served as the loading control. The bar chart shows quantification relative to controls. Data are presented as mean ± SEM, n = 3 mice per group; *P < 0.05.

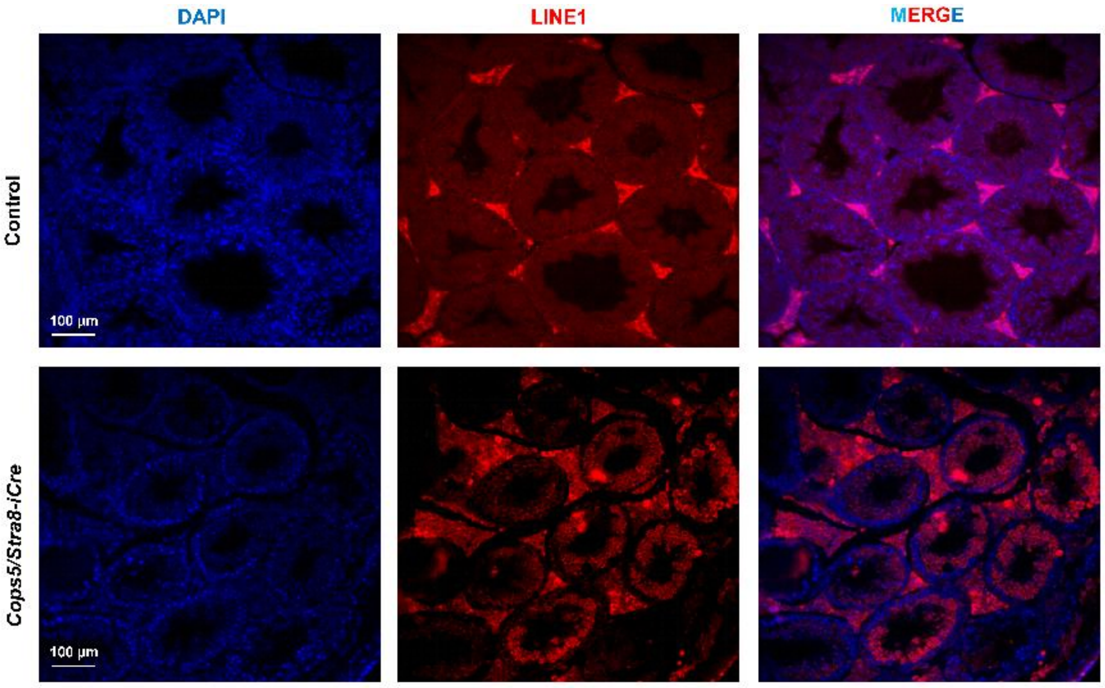

**Figure S4. Low-magnification immunofluorescence analysis of LINE1 in control and *Cops5* cKO testes.** Representative low-magnification images showing LINE1 immunoreactivity (red) in testis sections from control and *Cops5/Stra8-iCre* (cKO) mice; nuclei were counterstained with DAPI (blue). Scale bar, 100 μm.
